# Anuran Heart Metamorphosis: Anatomical Support for Pulmonary Blood Separation in the Early Aquatic Phase

**DOI:** 10.1101/2021.04.06.438645

**Authors:** Nina Kraus, Brian Metscher

## Abstract

1.

**Background:** In both larval and adult anurans, blood separation and respiratory physiology have remained an enigma. While various blood separation mechanisms have been proposed, the same structure is seen as playing a key role: the conus arteriosus. However, previous findings on its internal structure are contradictory, depending on the specifics of the 2D imaging methods used by different authors. To resolve this problem, we used high-resolution X-ray microtomography of whole *Bufo bufo* specimens to acquire the first detailed 3D descriptions of this complex structure through metamorphosis.

**Results:** In early tadpoles two small valvular openings develop at the ventricular-conal junction, providing two paths separated by the septum coni and continuing into the aortic arches. Thus, structures to support segregated pulmonary circulation are fully developed well before the lungs appear. The external gills undergo partial resorption and retreat asymmetrically into a gill chamber formed by a hyoidal cover, leaving only a single opening on the left side, the opercular spout.

**Conclusions:** The timing of events in *Bufo* circulatory development does not track the changing modes of respiration used by the developing tadpole. In particular, a system capable of double circulation carries only oxygen-depleted blood for a significant portion of the tadpole stage.

## 2. Introduction

Anuran heart function to this day presents a conundrum. While the gross adult heart morphology is relatively well understood, little is known about its morphology in detail, as well as about its function, physiology, and development. Most of our initial understanding of amphibian hearts has been an interpolation between the typical two-chambered fish heart with its single circulation and the post-embryonic four-chambered mammalian and avian hearts with their complete double circulation (Holmes 1975). This led to the initial conclusion that in the three-chambered anuran heart, oxygenated and deoxygenated blood mix and that this mixed blood supplies both the lungs and the rest of the body.

But judging vertebrate hearts based only on how much they resemble the mammalian one and understanding vertebrate hearts only as a continuum from fish to mammal disregards all the ways in which different morphologies benefit different ways of life. The amphibian heart is an unfortunate example for how traditional understanding of heart function neglects the particular adaptions of different heart morphologies. The amphibian heart is often dismissed as “less evolved” because its ontogeny seems to reflect some of the principal anatomical changes that occurred during vertebrates’ transition to land (Kolesová et al. 2007). However, it also has to be emphasized that there is no one “amphibian heart”. As with fish and reptilian heart morphology, amphibian hearts show significant intraclass variation, far more than avian or mammalian hearts do (Burggren 1988). This serves as a further indication that variations in heart morphology offer a source for unique adaptions to specific ways of life.

With the aim of understanding anuran heart function in more depth, further research reaching beyond the idea of simple mixing of oxygenated and deoxygenated blood was conducted starting in the 19th century. This led to two main opposing theories, and is illustrated in Fig. 1. It is well established that the amphibian heart generally consists of a sinus venosus leading into an atrium with a complete atrial septum, which connects to a ventricle with muscular trabeculae but no septation (Kardong 2009). Not having a septate ventricle is a unique condition within true lung-breathing vertebrates (Johansen & Hanson 1968). This ventricle further leads into a muscular conus arteriosus with two rows of valves and a septum, which is often called either spiral valve or septum coni. Even though the lack of a ventricular septum suggests either a complete mixing, or at most an imperfect separation of blood, it was soon assumed that there is at least some attempt to approximate the heart physiology of a complete double circulation (see de Graaf 1957).

**Figure 1:**
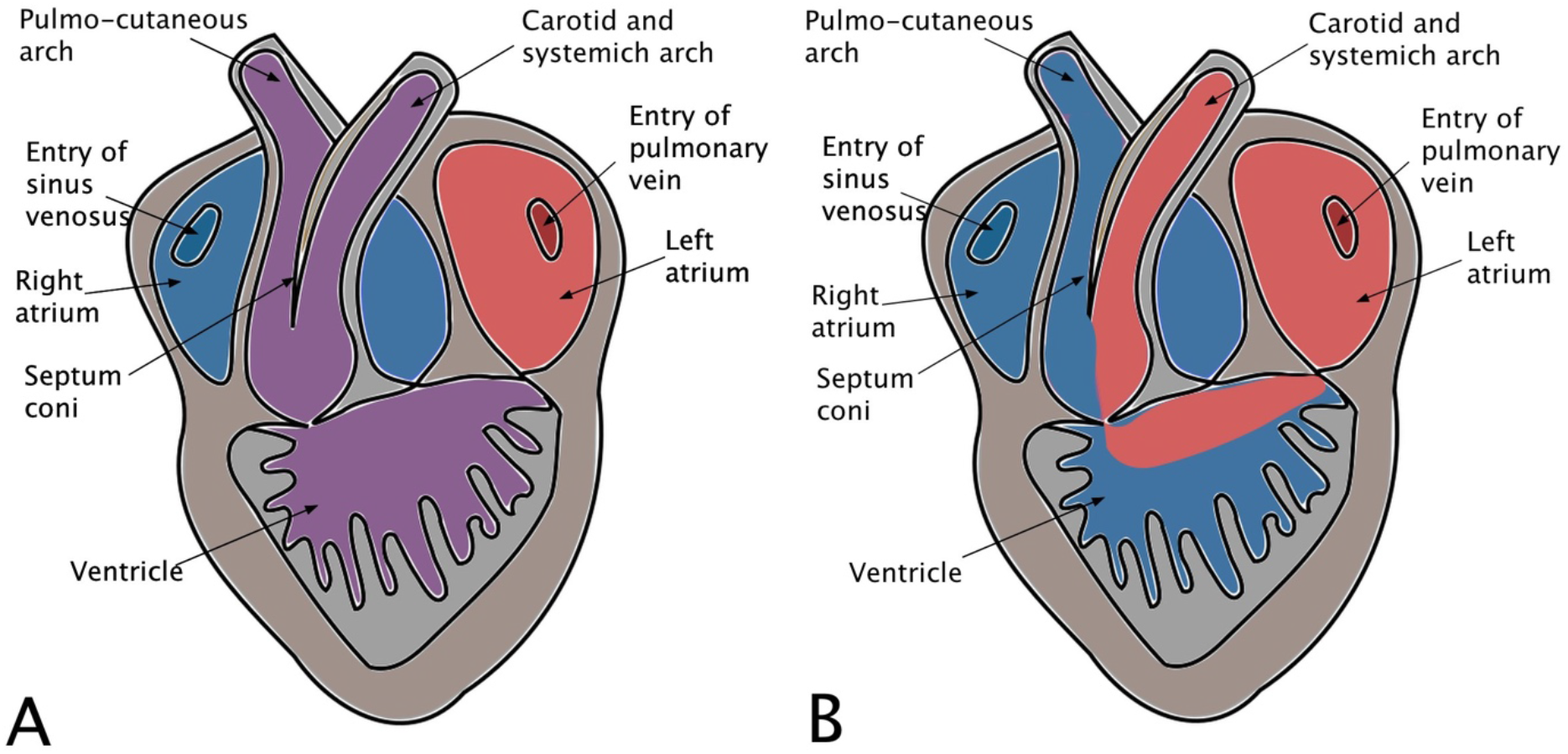
Two opposing theories about anuran heart function. A: Mixing Theory: In this model, deoxygenated blood from the body and oxygenated blood from the lungs simply mix in the ventricle and hence, stay mixed in their way through the conus and into the lungs and body. Here, the septum coni only aids in separating the blood into the aortic arches, not in keeping streams of blood of different oxygenation status separated. B: Classical Theory: Here, less oxygenated blood is said to be drawn into the ventricular meshwork and the ventricular systole sending it into the cavum pulmocutaneum and into the pulmo-cutaneous aortic arch. Later within the ventricular systole, the conus arteriosus is said to contract, causing the septum coni to contact with the conus wall at the base of the orifice at the junction between ventricle and conus, which closes of the cavum pulmocutaneum, so the more oxygenated blood can then enter the cavum aorticum and into the carotid and systemic aortic arch. Hence, the Classical Theory assumes an approximation of a functional double circulation.

This is described in the so-called Classical Theory, which is based on ideas by Brücke (1852) and was later expanded and modified by Sabatier (1873), as is explained in detail by de Graaf (1957). The Classical Theory (see Fig. 1B) describes a functional separation of oxygenated and deoxygenated blood by a combination of the atrial septum protruding into the atrio-ventricular opening, the drawing of the blood into the trabecular meshwork, and the septum coni continuing this separation of outflow blood throughout the conus arteriosus into the aortic arches. The last point is explained by only the pulmo-cutaneous aortic arch receiving the less oxygenated blood by a more favorable pressure gradient at the beginning of the ventricular systole. Then, at a later phase of the ventricular systole, the conus arteriosus contracts, which causes the septum coni to close off the opening into the cavum pulmocutaneum, so that oxygenated blood can only enter the cavum aorticum (see de Graaf 1957).

The opposing theory, the Mixing Theory (see Fig. 1A), claims that distribution of blood to the pulmo-cutaneous and systemic aortic arches is unregulated, meaning that there are separation mechanisms in play, but that they do not selectively work for oxygenated and deoxygenated blood. Instead they just split up mixed blood before it goes to the respective aortic arch (see de Graaf 1957).

The Classical Theory is based on some observations of the beating heart and theoretical considerations about its anatomy, as well as on injections and observations of different dyes such as ink, fluorescent dyes or x-ray-dense materials to see if and how separation occurs. The results showed at least partial separation of oxygenated and deoxygenated streams of blood (Acolat 1931, Morris 1974, Noble 1925, Simons 1957, Simons & Michaelis 1953).

Proponents of the Mixing Theory however, tried to replicate the injection experiments but failed to get the same results (e.g. Foxon 1947, 1951, 1953). However, it has to be pointed out that this could easily be because of minute details of experimental design. For example, when working with dyes the thick muscular walls of the ventricle and the conus arteriosus make it fairly difficult to observe the distribution of the dye and close to impossible to observe anything behind the septum coni relating to the area of the cavum pulmocutaneum. Additionally, great care is required when deciding on an adequate amount of dye or similar material for such an experiment. While using too little agent would make the tracing of blood flow impossible, using too much could potentially increase the blood volume to such a degree that potential separating mechanisms would lose their functionality. And of course, the simple act of opening the animals’ chest and its pericardium are already invasive and could already influence the results (see de Graaf 1957).

In short, some researchers argued that the anuran heart’s anatomy enables it to form a functional double circuit and to keep different kinds of blood almost completely separated, while others argued for random mixing and distribution of oxygenated and deoxygenated blood. Still others thought of it as more of a partial separation, even arguing that anurans are able to adjust the degree of mixing based on physiological conditions such as temperature, and hence oxygen content of the water (Hillman et al. 2014, Simons & Michaelis 1953). Overall, there is no clear answer as to the exact mechanism and degree of potential separation of oxygenated and deoxygenated blood. Thus, it is still under discussion how the anuran heart manages to control and organize blood flow to and from its different sites of gas exchange.

What is known is that the amphibian cardiovascular system uses a single ventricle to generate pressure that drives blood through the conus arteriosus and into parallel systemic, carotid, and pulmo-cutaneous circuits (Wang et al. 1999). Oxygenated blood then returns through the left atrium and systemic venous blood through the right one. This parallel vascular arrangement would, as supporters of the Classical Theory pointed out, at least theoretically have the potential to act as a functional double circulation (Acolat 1931, Morris 1974, Noble 1925, Simons 1957, Simons & Michaelis 1953), as well as potentially even enable the delivery of more or less blood to the respiratory surfaces depending on metabolic state (Hillman et al. 2014, Simons & Michaelis 1953). However, few details are known about respiration physiology, since studying anuran heart function proves to be difficult.

The need to acknowledge amphibian heart morphology as a unique specialization becomes especially apparent when looking at anuran larvae. Even less is known about their respiratory physiology than about their adult counterparts. Tadpoles cannot be understood merely as an extension of the embryonic phase but are a highly specialized life form having unique traits within the anuran life cycle (McDiarmid & Altig 1999). Especially the respiratory system undergoes profound morphological changes throughout metamorphosis: the transition from water to land is usually accompanied by a change not only in the way their respiratory surface is ventilated, but also in the respiratory surface itself.

Early tadpoles start out relying predominantly on cutaneous gas exchange across the highly vascularized skin with additional small external gills containing only a few capillary loops (McIndoe & Smith 1984). Unlike in salamanders, in anuran larvae those external gills are soon replaced with more efficient filamentous internal gills on four gill arches which are ventilated via a unidirectional flow of water leaving the pharyngeal cavity through a single opercular spout on the animal’s left side. As adults, pulmonary ventilation is the main mode of oxygen uptake, still accompanied by cutaneous respiration, especially for discharging CO_2_ (Burggren & Infantino 1994, Feder & Burggren 1985). However, cutaneous breathing seems to account for a larger percentage of gas exchange than the active ventilation of gills (Burggren & West 1982).

This transition between different modes of breathing is not abrupt but progressive. Lung ventilation can, depending on species, start at any time up to metamorphic climax once the gills have already started to degenerate, but the two modes are, at least for a short period of time, used simultaneously along with cutaneous breathing (Burggren & Infantino 1994, Burggren & Pinder 1991, Feder 1984).

Additionally, the morphological basis for the transition from external to internal gills in anurans has yet to be described in detail. A thorough literature search turned up only a short note by Rugh (1951) saying that the hyoid arch goes on to form parts of the tadpoles’ operculum. Another mystery yet to be solved is the way in which tadpoles ventilate their lungs. Along with being interesting questions by themselves, knowing more about breathing modes and their morphological basis in tadpoles will aid in resolving how the anuran heart functions both in larval and adult animals.

Nonetheless, it is still under discussion how the anuran heart manages to control and organize blood flow to and from all different sites of gas exchange, especially simultaneously. Thus, in both anuran larvae and adults, blood separation and respiratory physiology have remained an enigma. But while different mechanisms of blood separation have been proposed by supporters of both the Mixing Theory and of the Classical Theory, both agree on the same structure as playing an important role in blood separation in anurans: the conus arteriosus (Sharma 1961, Simons 1957).

In the form found in amphibians, the conus arteriosus is unique among tetrapods. In amphibians, as well as in lungfish, the conus is a prominent structure, connecting the ventricle with the truncus arteriosus (Johansen & Hanson 1968). In amniotes, the conus arteriosus exists embryonically where it is usually referred to as the bulbus cordis. During development, this bulbus cordis then becomes incorporated into the bases of the aorta and the pulmonary artery, where it contains the semilunar valves. These control the emptying of blood from the ventricle into the truncus arteriosus, which later in development constitutes the aortic arches (Holmes 1975, Kardong 2009).

What distinguishes the anuran conus from the amniote one (other than size) is the number of rows of valves within it. While amniotes only have one row of valves, amphibians have two, one at the bottom of the conus and one at the top (Holmes 1975, Kardong 2009). Especially considering non-teleost coni with their several rows of valves, this pattern seems to suggest that the condition found in anurans to be an ancestral or intermediate state for tetrapods. However, lungfish show the same condition as anurans do, suggesting that this condition is based more on adaption than on phylogeny. Additionally, anurans (and lungfish) have a prominent structure between those rows of valves, the septum coni, which separates the anuran conus into two cava, the cavum aorticum and the cavum pulmocutaneum, and is said to play a crucial role in the separation of blood in anurans (Johansen & Hanson 1968).

Not only are the exact function and mechanism of the conus arteriosus still under discussion, but even its general morphology is not agreed upon. Comparison of published descriptions of the conus arteriosus reveals important contradictions regarding the morphology of the valves within the conus arteriosus, and regarding the spiraling pattern of the septum coni and its anterior attachment to the wall of the conus arteriosus (Ison 1967, 1968, Sharma 1961, Simons 1957).

The number of valves reported within the conus arteriosus varies dramatically depending on whether the authors sectioned their samples horizontally or transversely, as pointed out by Ison (1967). To try to solve this problem, Ison produced and analyzed both transversal and horizontal histological sections, clearly trying to get a more three-dimensional picture of conus morphology. However, due to the technical and methodological limitations of his time, Ison’s results are still hard to interpret.

Following conus morphology throughout development would also enable better understanding of anuran hearts. Ison (1968) tried this as well, but again, histological sectioning limited his results, since this time-consuming method did not enable him to realistically cover the extremely rapidly developing anuran heart fully. Other than Ison’s work, most developmental studies focused on very early heart development (e.g. Mercola et al. 2010), stopping shortly before the main parts of the heart are fully differentiated, or the work was simply not focused on conus morphology (Kolker et al. 2000).

In early anuran heart development, the typical three-layered organization of early vertebrate hearts also has yet to be considered. Embryonic vertebrate hearts consist of an external layer, the myocardium, which is made up of two layers of cells. On the inside of the heart sits a single-cell layer of endocardium. In between those two layers lies cardiac jelly, an extracellular matrix containing collagen fibers. During development, this layer of cardiac jelly gets either displaced by endocardial cells growing towards the myocardium or is replaced by endocardial cells migrating into it, with the exact mechanism depending on the precise area within the heart (Courchaine et al. 2018, Kirby 2007). This phenomenon should be important for adult heart morphology, since the way cardiac jelly gets replaced by cardiac cells is coordinated via hemodynamic influences and thereby influences the formation of structures within the heart, including valves and septa. While the existence of this enigmatic extracellular matrix was already acknowledged before most anuran heart morphology research was conducted (Davis 1924), most authors seem to not have been aware of it and its peculiar properties. Considering the cardiac jelly in the context of anuran heart development could reveal further details about the heart morphology found in adult specimens by following the structures throughout the larval phase.

To fill in these gaps in the literature and to resolve the actual morphology of the conus arteriosus, we used 3D micro-CT imaging to analyze the structure of the anuran conus arteriosus throughout metamorphosis in order to establish a strong foundation on which to base future research on the anuran cardiovascular system. Additionally, we described gill morphology during development to further our understanding of the transition between the different modes of respiration in larval anurans.

## 3. Methods

### 3.1. Samples

Specimens of *Bufo bufo* were acquired from the herpetological collection of the Natural History Museum of Vienna. Samples were previously fixed and stored in a solution of 50% EtOH and 5% formalin. Using a stereomicroscope, developmental stage was assigned according to the Gosner staging table (Gosner 1960), which is based on external morphological characteristics of anuran larvae. A list of the specimens appears in the appendix/supplementary material.

### 3.2. Staining

Specimens younger than Gosner stage 43 were stained using 0.3% PTA in 70% EtOH (Metscher 2009). First, those specimens were washed in 70% EtOH three times, for about 15 minutes each. Then, those specimens were transferred to the staining solution for 24-48 hours, depending on size. Subsequently, the specimens were washed again in 70% EtOH for at least 1 hour with one change to fresh 70% EtOH during this period.

Due to the increased thickness of the epidermis in older specimens, starting from Gosner stage 43, aqueous iodine (Lugol’s solution, IKI; (Metscher 2009) was used as a contrast stain. These specimens were initially rehydrated by a descending alcohol series (50% and 30% EtOH for 30-60 minutes each). Later, those specimens were washed in distilled water twice for at least 15 minutes each. After rehydration, the specimens were moved to IKI solution for about 48 hours and after that were again washed in dH_2_O twice, for about 30 minutes each.

Additionally, a single specimen was stained with Rose Bengal (an iodated eosin analog; Chroma 1A 182) at 0.5% in 50% EtOH, and after micro-CT scanning, it was embedded in paraffin and sectioned at 7μm. The slides were dewaxed in two washes of xylene (5 minutes each) and then cover slipped without further staining.

### 3.3. Micro-CT imaging

After staining, all specimens were mounted in 1% aqueous agarose (Metscher 2011) for scanning. Specimens were scanned using a Xradia MicroXCT microtomography imaging system (Zeiss X-Ray Microscopy) with a tungsten x-ray source operated at 40-60 kV and 4 W. Individual projection images were taken every 0.20° over a 180° scan, with 40 s exposure times. The tomographic sections were reconstructed with voxel sizes of 0.89– 3.99 μm using the XMReconstructor software supplied with the scanner.

### 3.4. Analysis

Scans were conducted starting from Gosner stage 18, but the main focus was on scans starting from Gosner stage 19 (the time the heart starts to beat) up to stage 46 (a small froglet that has just finished metamorphosis). Earlier development was excluded from this study, since it has already been covered extensively by other authors (e.g. Rugh 1951).

Initially, one specimen was scanned per developmental stage. If initial analysis showed that developmental changes are too rapid to be fully covered by the Gosner table, which is only based on external morphological characteristic, more scans were conducted of the specific developmental stage.

Obtained images of specimens were examined and described using the Xradia XM3DViewer software, Dragonfly 3.6 (http://www.theobjects.com/dragonfly) and Amira 6.4 or 2020.2 (ThermoFisher Scientific). Segmentations were made using the Dragonfly 3.6 segmentation tool. Images and segmentation where additionally compiled and used for making schematic drawings of the overall heart morphology, as well as conus morphology. Those drawing were digitalized using Inkscape (http://www.inkscape.org/). General image manipulation and labeling were done with Fiji and all images were contrast enhanced (Schindelin et al. 2012) and its plugin FigureJ (Mutterer & Zinck 2013).

## 4. Results

### 4.1. Tadpole Heart and Outflow Tract Development

At Gosner stage 18, the two mesodermal heart tubes are already fully fused, the heart has already started to coil into its distinct S-shape, the main regions of the heart can already be distinguished.

At Gosner stage 19 (Fig. 2) the heart tubes’ inflow area (future atria) is still connected to the yolk via the vitelline veins. With the whole heart coming to lie in the animals’ center line the inflow area is bent to the right and lies dorsally under the mouth opening. The heart tube then bends ventrally and to the left, leading into the ventricle. The ventricle itself already started to balloon at this point, taking on a wide U-shape, with its most dorsal curvature being concave. Subsequently, the heart tube twists dorsally and to the right, leading into the outflow area (the future conus).

**Figure 2:**
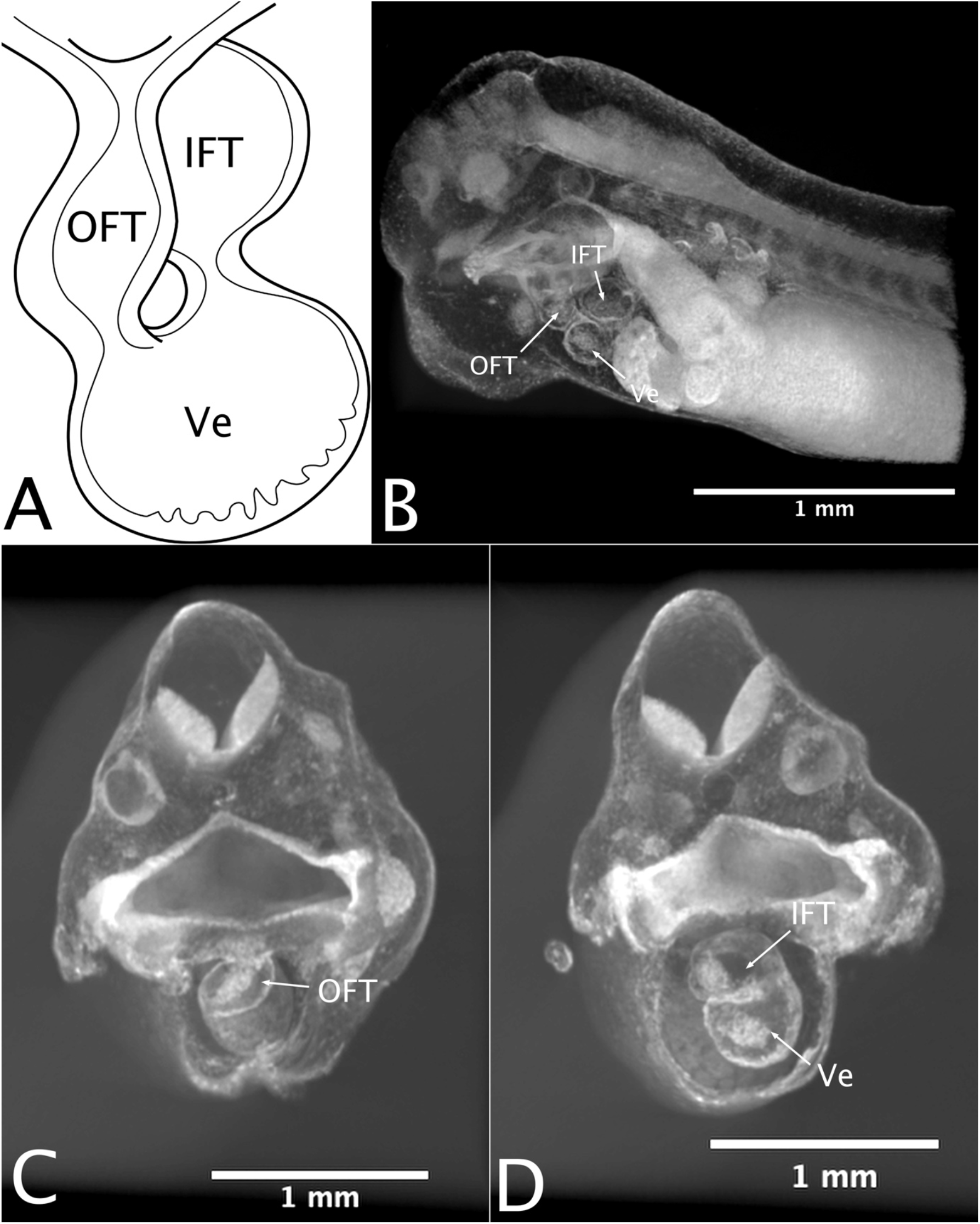
Anuran heart morphology at Gosner 19. A: Schematic drawing of the gross heart morphology at this developmental stage from the ventral view. B: lateral view of a virtual sagittal cutaway. C: rostral view of a virtual transversal cutaway. D: rostral view of a virtual transversal cutaway, more posterior than C. IFT inflow tract, OFT outflow tract, Ve Ventricle.

At this point in development, the formation of trabeculae within the ventricle begins, starting from the structures middle, the apex, and proceeding towards the inflow and the outflow area. In those areas themselves, however, the embryonic heart tissues have not yet started to differentiate morphologically, so the typical three-layered structure of the embryonic vertebrate heart is still fully present.

Little changes up to Gosner stage 21, when the rate of heart development starts to increase. Hence, developmental changes are too rapid to be fully covered by the Gosner table, so several scans were made of specimens within this stage.

During stage 21 (Fig. 3), the inflow area/atrium starts to balloon and increases in diameter. The ventricle continues to balloon as well, and thereby becomes rounder and less U-shaped than previously, meaning the initially concave top curvature becomes more convex. Trabeculation proceeds towards outflow and inflow area, so that the full width of the ventricle is in the process of becoming trabeculated. Already existing trabecula become more pronounced. At the top curvature are no trabecula, so cardiac jelly remains. The outflow area becomes more elongated and tilts more ventrally and less to the right than prior. Its layer of cardiac jelly becomes thinner.

**Figure 3:**
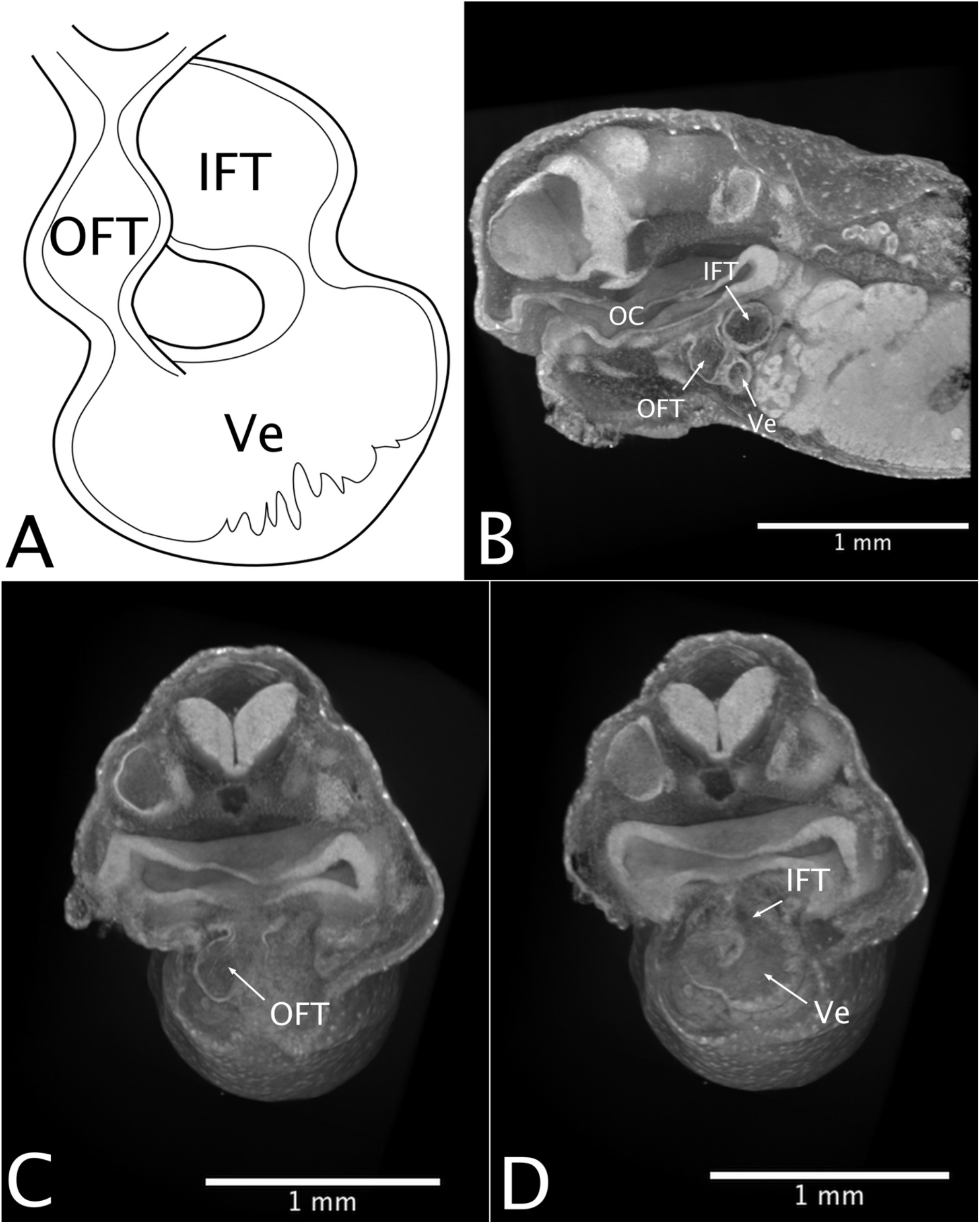
Anuran heart morphology at Gosner 21. A: Schematic drawing of the gross heart morphology at this developmental stage from the ventral view. B: lateral view of a virtual sagittal cutaway. C: rostral view of a virtual transversal cutaway. D: rostral view of a virtual transversal cutaway, more posterior than C. IFT inflow tract, OFT outflow tract, Ve Ventricle, OC oral cavity.

With stage 22, the S-shaped coiling of the heart tube becomes tighter, so the distances between outflow area, ventricle and inflow area become smaller and the outflow area comes to lie in front of the inflow area (see Fig. 6 for relation of those structures to each other). The ventricle is already very round and appears fully trabeculated.

Starting at Gosner stage 22 the atrial septum starts to appear. Also, cells of the endocardial layer within the conus start to migrate into its cardiac jelly. This leads to a small, tongue-shaped structure to protrude from the right dorsal side of the conus reaching downwards and curving in the same fashion as the conus itself does (Fig. 4B and 6). This protrusion will go on to form the septum coni. It initially contacts a ring of tissue at the junction of the conus arteriosus and the ventricle, but at a later phase of this Gosner stage, only its tip still contacts the valves and the septum coni elongates. At this point however, there is only a single cavum within the conus, through which blood could potentially flow.

**Figure 4:**
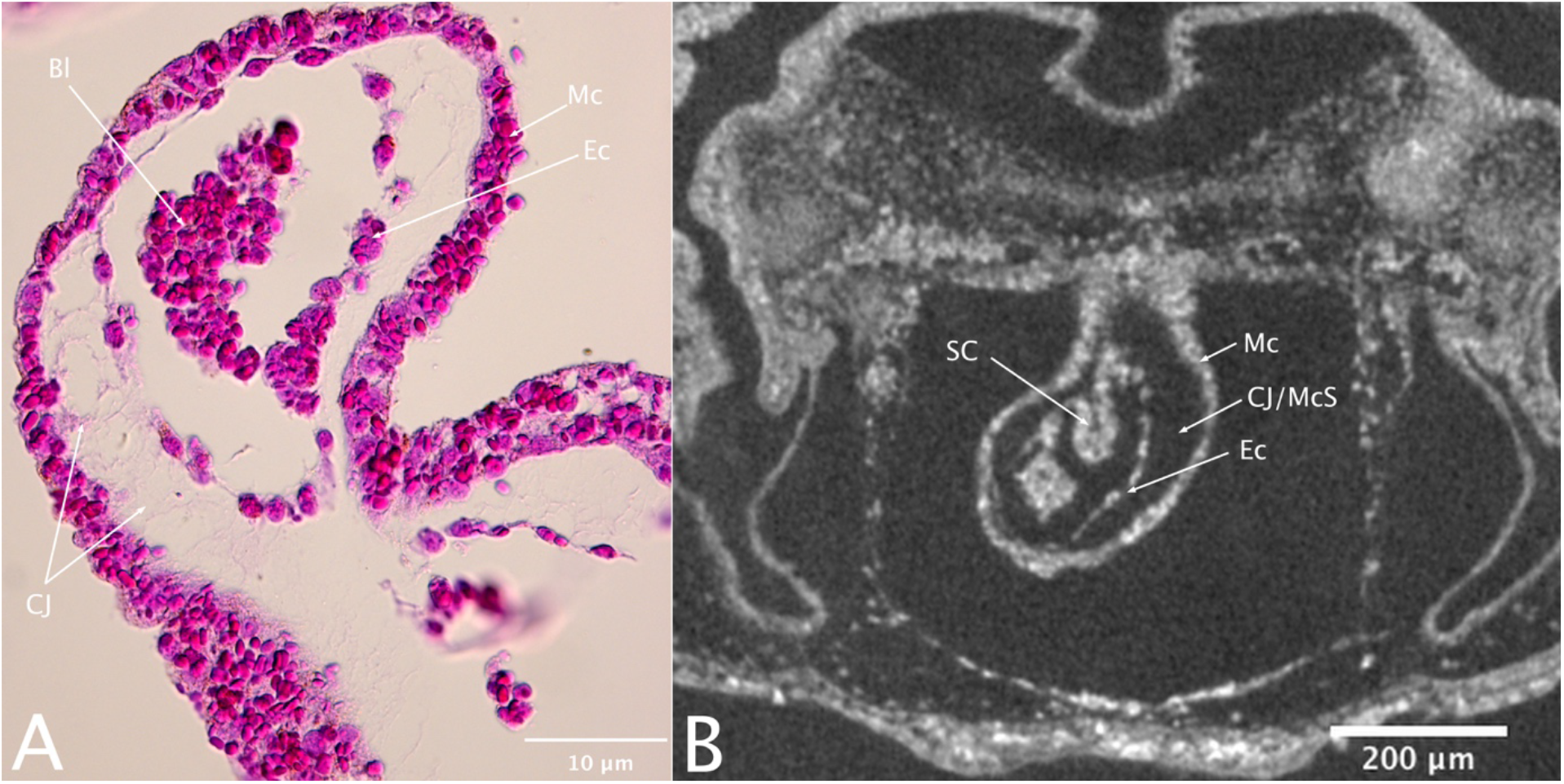
The typical three-layered structure of an embryonic vertebrate heart with the myocardium on the outside and endocardium on the inside of the heart with the cardiac jelly in between. A: Histological section through the conus arteriosus of a tadpole at Gosner stage 22. B: Virtual section through the conus arteriosus of the same specimen (more ventral) showing the formation of the septum coni. Bl Blood, CJ Cardiac jelly, CJ/McS Cardiac jelly/Myoendocardial space, Ec Endocardium, Mc Myocardium.

The ring of tissue at the bottom of the conus, as well as on its top start to form from endocardiac cushions that start to appear at Gosner stage 23 (Fig. 5). The bottom ring of tissue develops a small protrusion within Gosner stage 23 that contacts the septum coni. The septum coni itself starts to grow thicker and to become more prominent and looks strongly nucleated (see Fig. 7). At this stage as well, a small extra opening from the ventricle into the conus starts to appear in this ring of tissue. It lies more caudally than the main opening and remains rather small and narrow (see Fig 11B).

**Figure 5:**
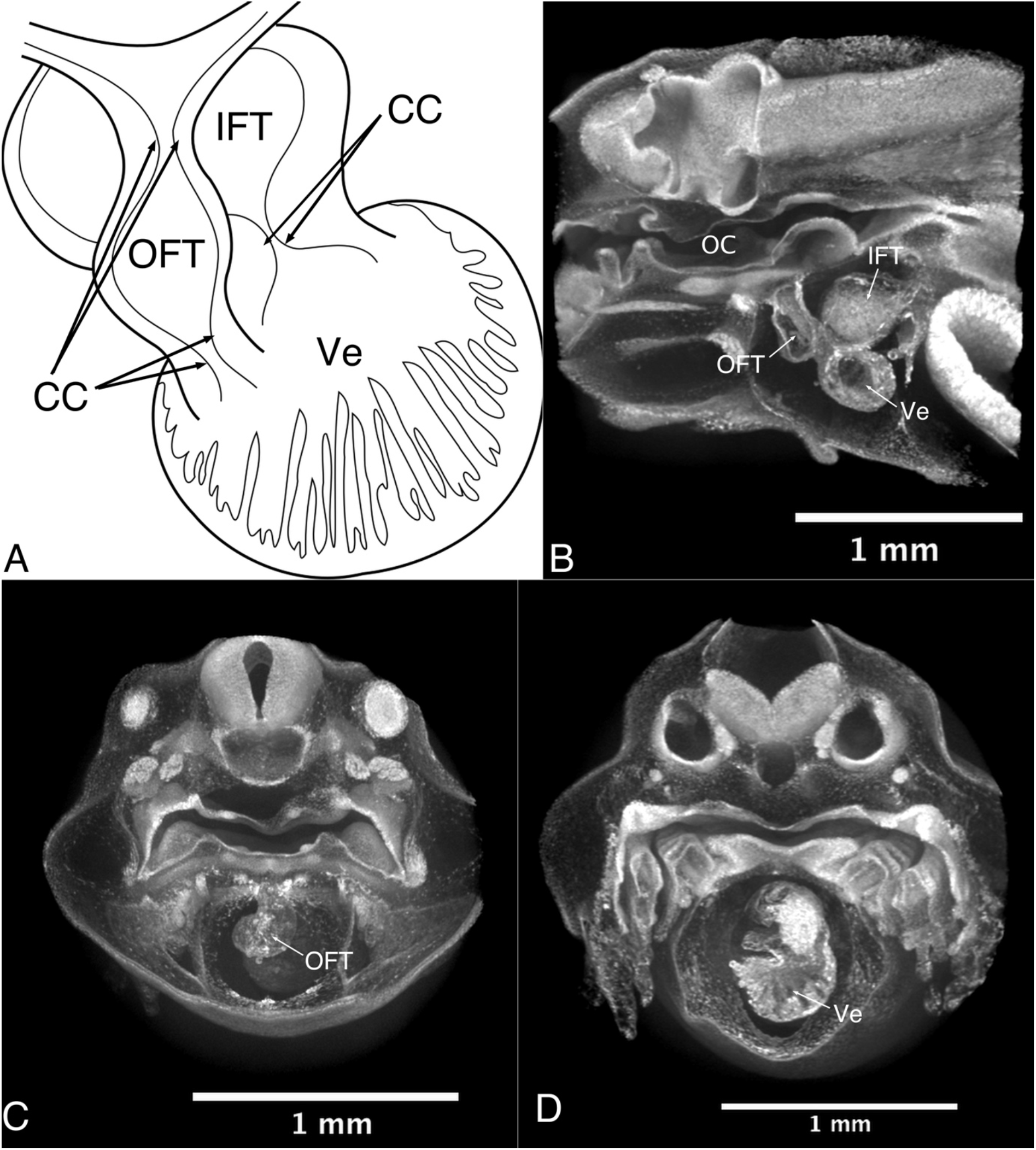
Anuran heart morphology at Gosner 23. A: Schematic drawing of the gross heart morphology at this developmental stage from the ventral view. B: lateral view of a virtual sagittal cutaway. C: rostral view of a virtual transversal cutaway. D: rostral view of a virtual transversal cutaway, more posterior than C. CC cardiac cushions, IFT inflow tract, OC oral cavity, OFT outflow tract, Ve Ventricle.

**Figure 6:**
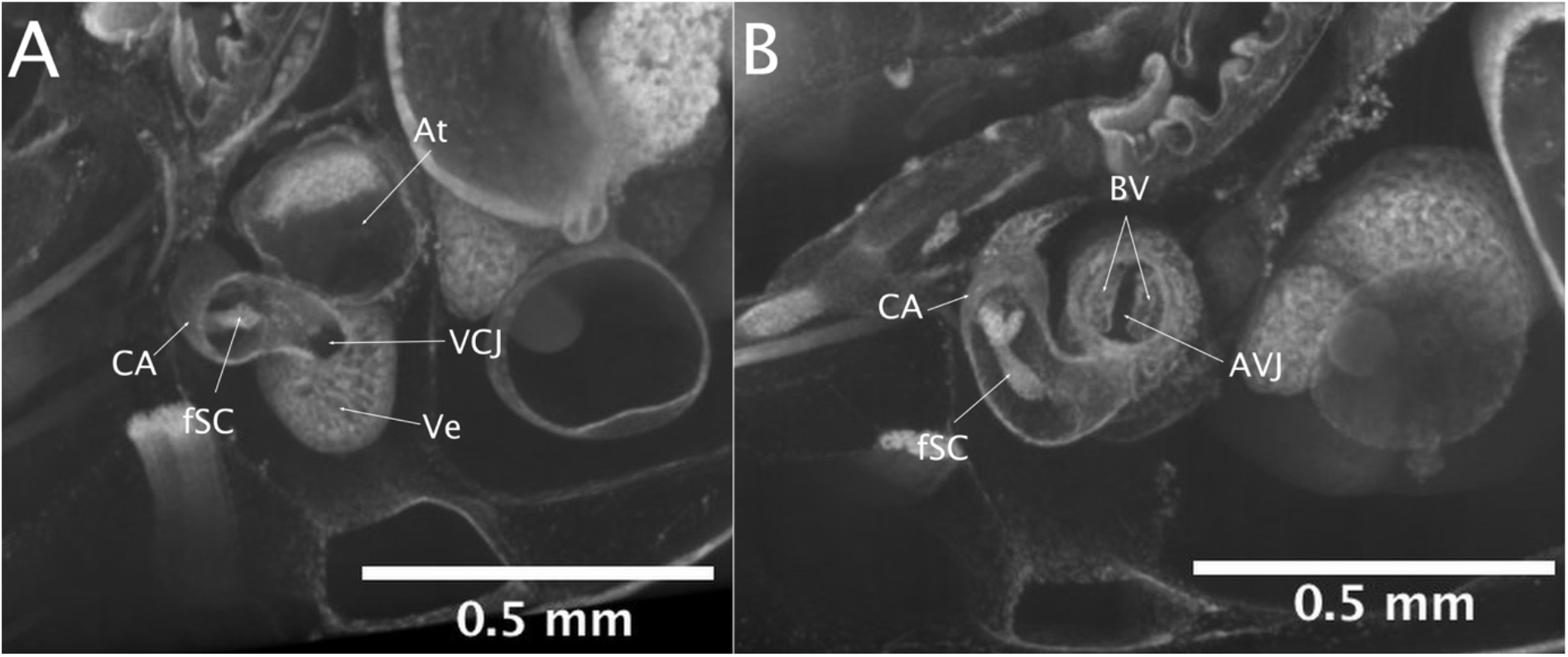
Virtual cutaway of a 3D reconstruction of a tadpole at Gosner stage 25 showing how the atrium mouths into the ventricle at the atrio-ventricular juncton and the bicuspid valves of the atrium, and where the conus mouths into the ventricle at the ventricular-conal junction, as well as as the spiraling pattern of the conus and the septum coni. At atrium, AVJ atrio-ventricular junction, BV bicuspid valves, CA conus arteriosus, fSC forming septum coni, VCJ ventricular-conal junction Ve ventricle.

**Figure 7:**
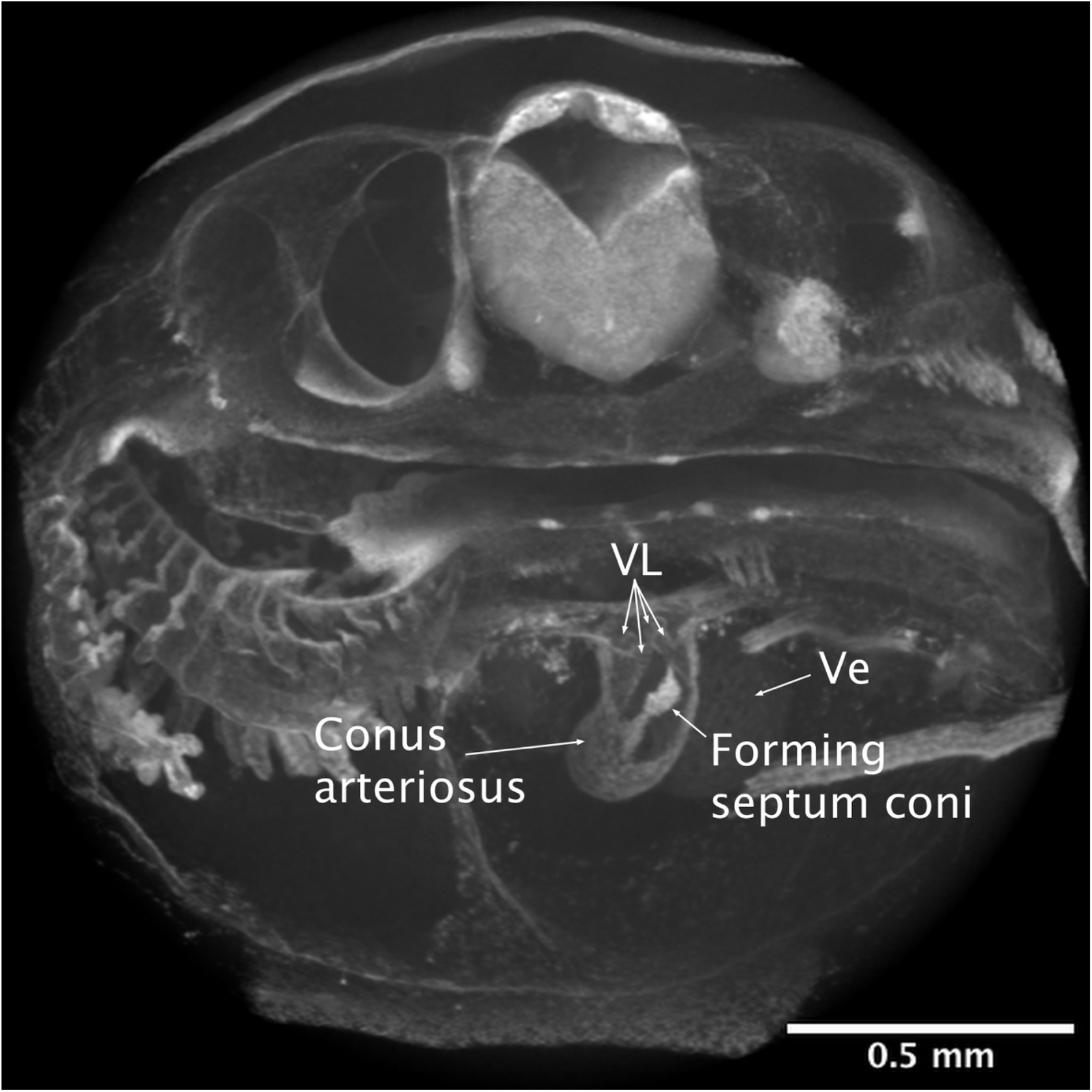
Virtual cutaway through the transverse plane of a 3D reconstruction of a tadpole at Gosner stage 24 showing the now distinct four leaflets of the two top valves and the formation of the septum coni which contacts its tongue shaped protrusion. Note the higher overall X-ray density of the septum coni, owing to its high density of nuclei. Ve ventricle, VL valve leaflets.

From Gosner stage 24 up to 25, the endocardiac cushions at the top (Fig. 7) and bottom (Fig. 13) of the conus develop into defined valves which at this point are all connected to other structures within the conus via a thin lining of a sheet of tissue (as is shown in Fig 11A). Once the bottom valves are more defined, it is visible that the septum coni contacts the smallest of those valves (the left one), which lies closest to the atrial opening of the ventricle. Additionally, those valves are connected to the septum coni not only by this protrusion, but also by a lining of tissue of the right side of the conus. On the left side, this lining is significantly thinner. While there seemed to be minor individual differences, this bottom ring of valves consists of two main valves and a small extra protrusion that contacts the septum coni, which can be understood as its own small valve (see Fig. 13). At the top there are two main valves, each made up of two leaflets, but again, they form a continuous ring of tissue that is connected to the conus wall (see Fig. 7).

At this point in heart development, the septum coni grows an additional small protrusion that grows upwards and reaches into the middle of the top row of valves (see Fig. 9, 11B and 12). This leads to those structures forming a “Y” that divides three small chambers within the junction of the conus into the aortic arches (see Fig 8, 9, 11B, and 12). The dorsal one of those is connected to the pulmo-cutaneous arch, while the other two, more ventrally lying cava are connected to the systemic and the carotid arches. The reduction of the conus lining and the connection of the tongue-shaped septum coni to the bottom valves, as well as the appearance of the small extra opening at the base of the conus results in two cava within the conus, through which blood could potentially flow. Additionally, the small protrusion on one of the bottom valves of the conus has lost direct contact to the conus wall, creating a very small, additional way blood could potentially enter the conus from the ventricle. Since this protrusion is connected to the septum coni, this opening aligns with the left cavum of the conus, the cavum pulmocutaneum, while the large opening formed by the ring of valves aligns with the right cavum, the cavum aorticum. The right cavum connects to the openings of the two valves into the systemic and carotid arches, and the cavum on the left side connects to the opening to the pulmo-cutaneous arches. From this point on there are two cava within the conus, that would support two fully separated streams of blood throughout the whole length of the conus and its junction into the aortic arches (see Fig. 12).

**Figure 8:**
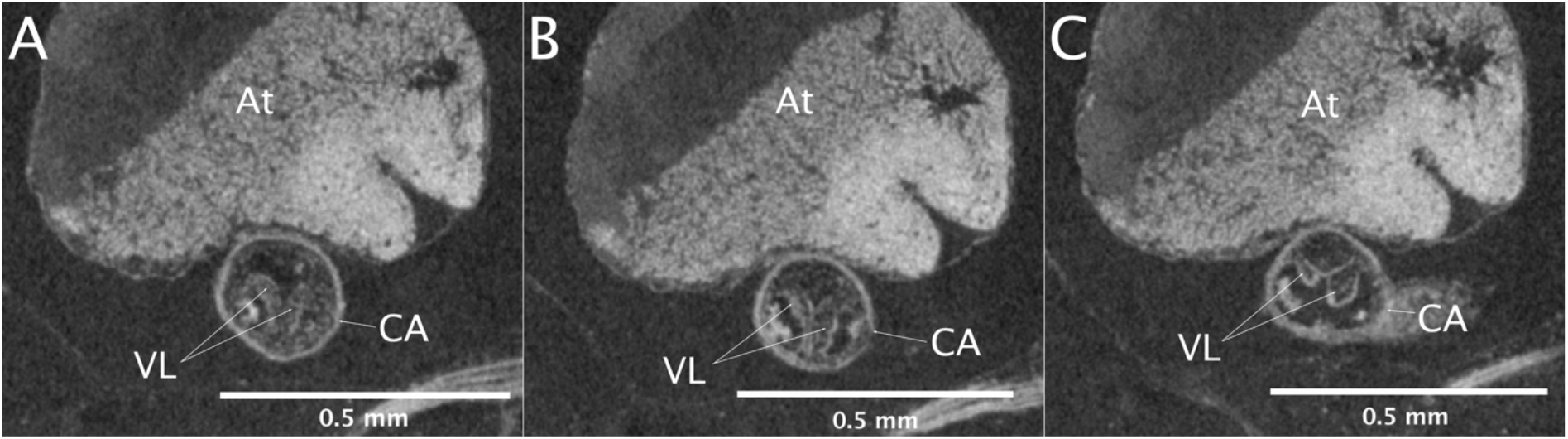
Virtual section through the conus arteriosus of a tadpole at Gosner stage 29 showing the leaflets of the top valves contacting each other to form the distictive Y-shape, A being the most caudal one, C the most rostral one. At Atrium, CA Conus arteriosus, VL Valve leaflets.

**Figure 9:**
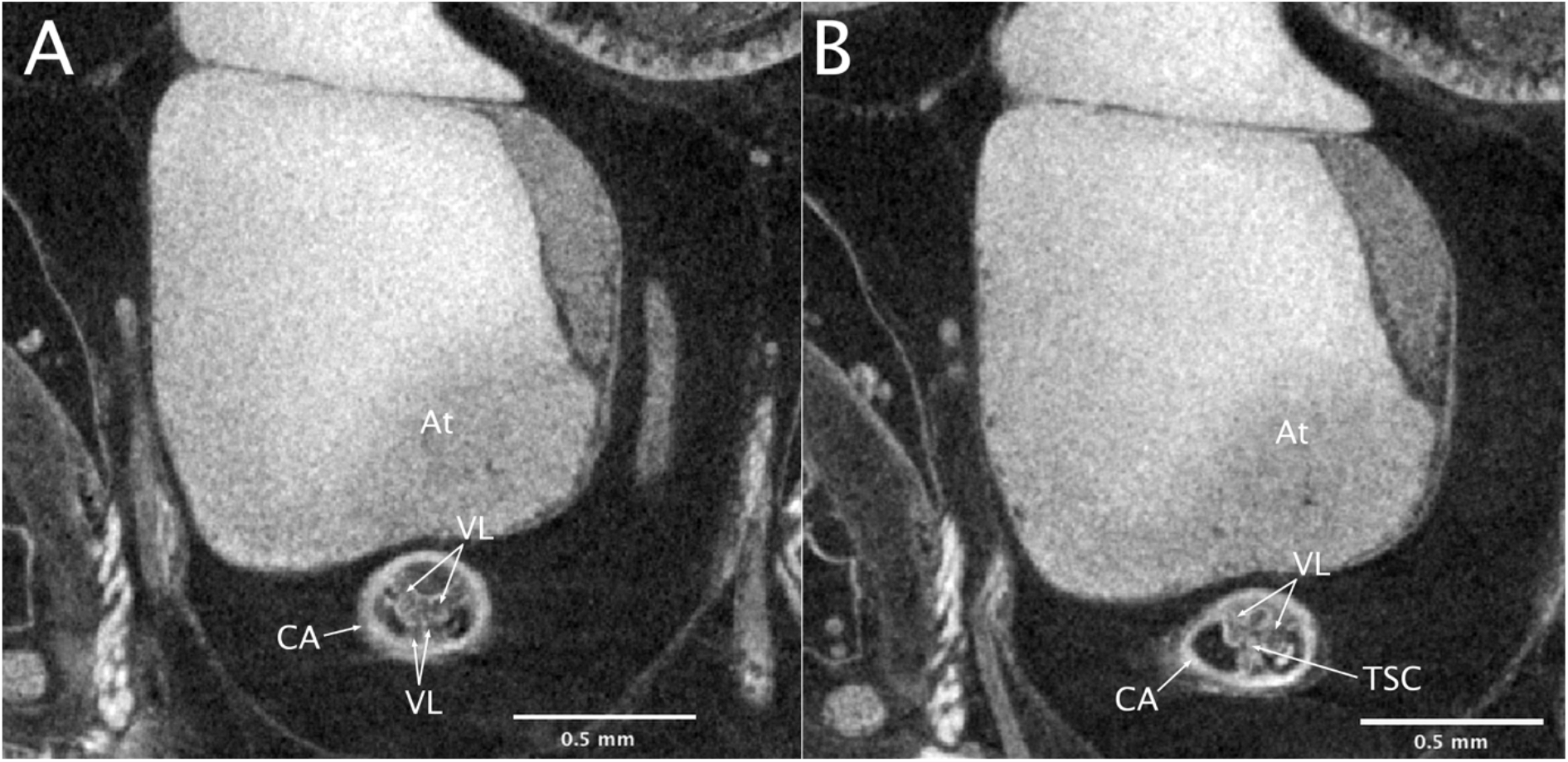
Virtual section through the conus arteriosus of a tadpole at Gosner stage 36 showing the leaflets of the top valves contacting each other to form the distictive Y-shape. Note that the more dorsal valve leaflets closing off the entry of the cavum pulmocutaneum into the pulmo-cutaneus aortic arch reach further rostral (B) than the ones closing off the cavum aorticum and reaching into the systemic and carotid aortic arches. At Atrium, CA Conus arteriosus, TSC Tip of septum coni, VL Valve leaflets.

**Figure 10:**
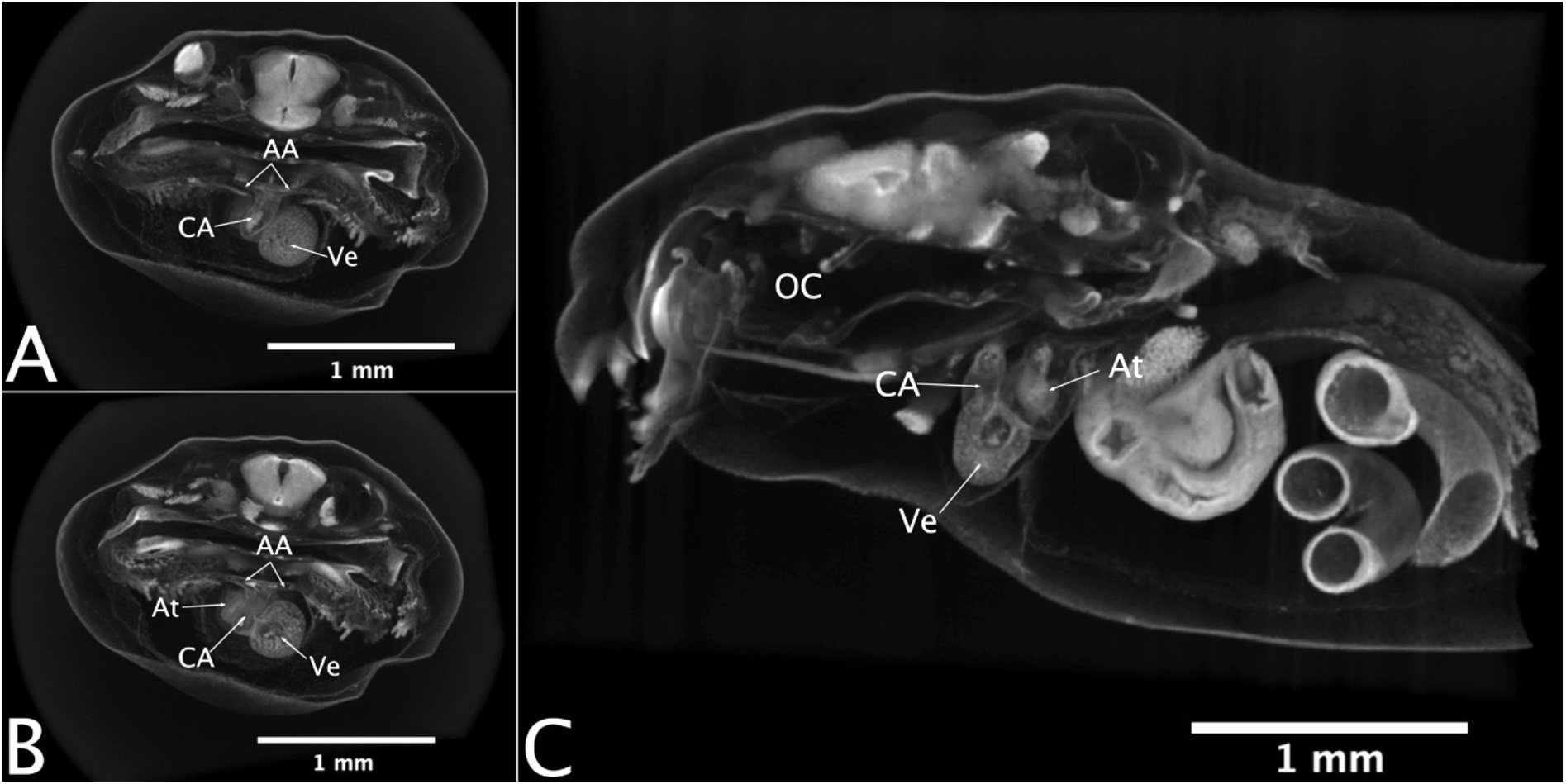
Anuran heart morphology at Gosner 25. A: rostral view of a virtual transversal cutaway. B: rostral view of a virtual transversal cutaway, more posterior than A. C: lateral view of a virtual sagittal cutaway. AA aortic arches, At Atrium, CA conus arteriosus, OC Oral cavity, Ve Ventricle.

**Figure 11:**
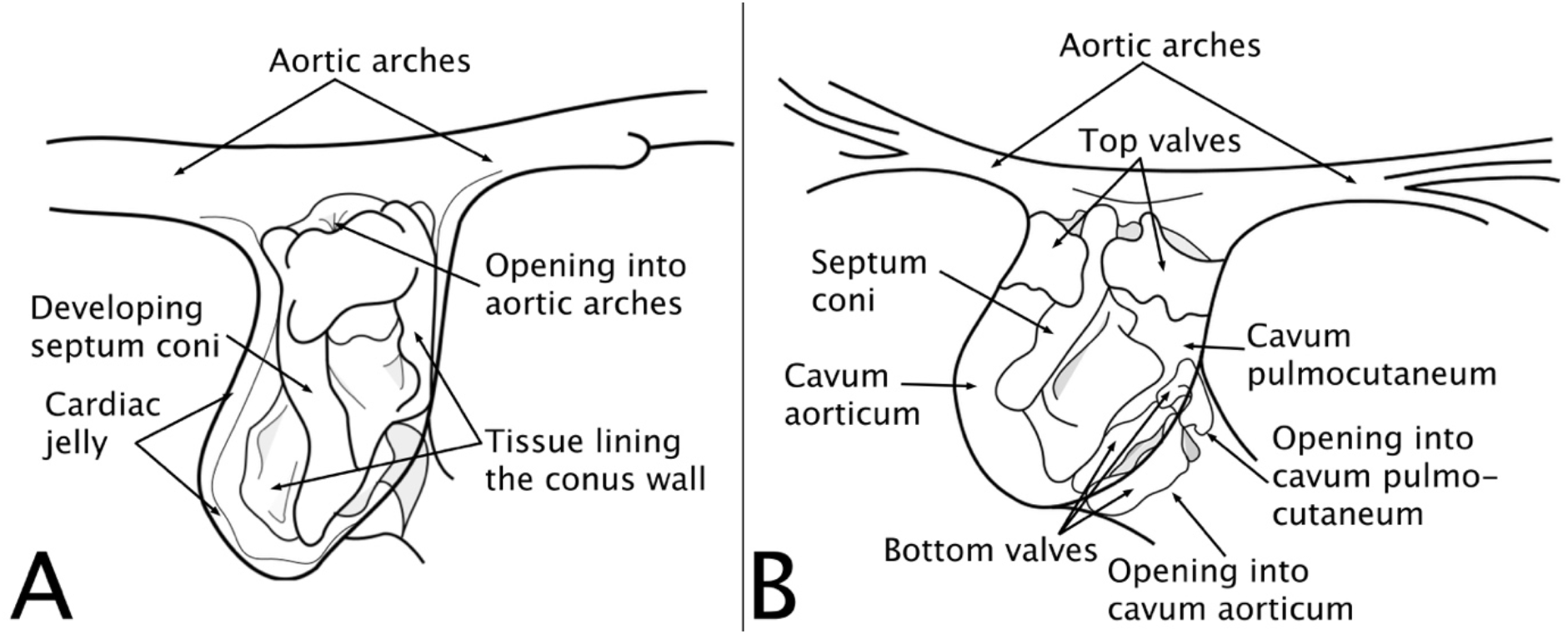
Schematic drawing of the ventral view of the morphology of the conus arteriosus through anuran heart development. A: Conus arteriosus at Gosner stage 22: Here, the septum coni does not yet lie close to rostral wall of the conus, so instead of dividing it into two chambers, it allows for blood flow in front of it. At this point in development, the conus shows no distinct valves and is still mostly lined by thin sheet of tissue. B: Conus arteriosus at Gosner stage 43: The septum coni has already grown significantly towards the rostral conus wall, but is not in direct contact with it. The two main bottom walls and the small extra one/extra leaflet are already fully formed, allowing for two separate entries for blood into the conus. The morphology suggests that blood entering via the small opening at the ventricular-conal junction will flow into the cavum pulmocutaneum, while blood entering via the main opening leads into the cavum aorticum. The two cava are separated via the septum coni.

**Figure 12:**
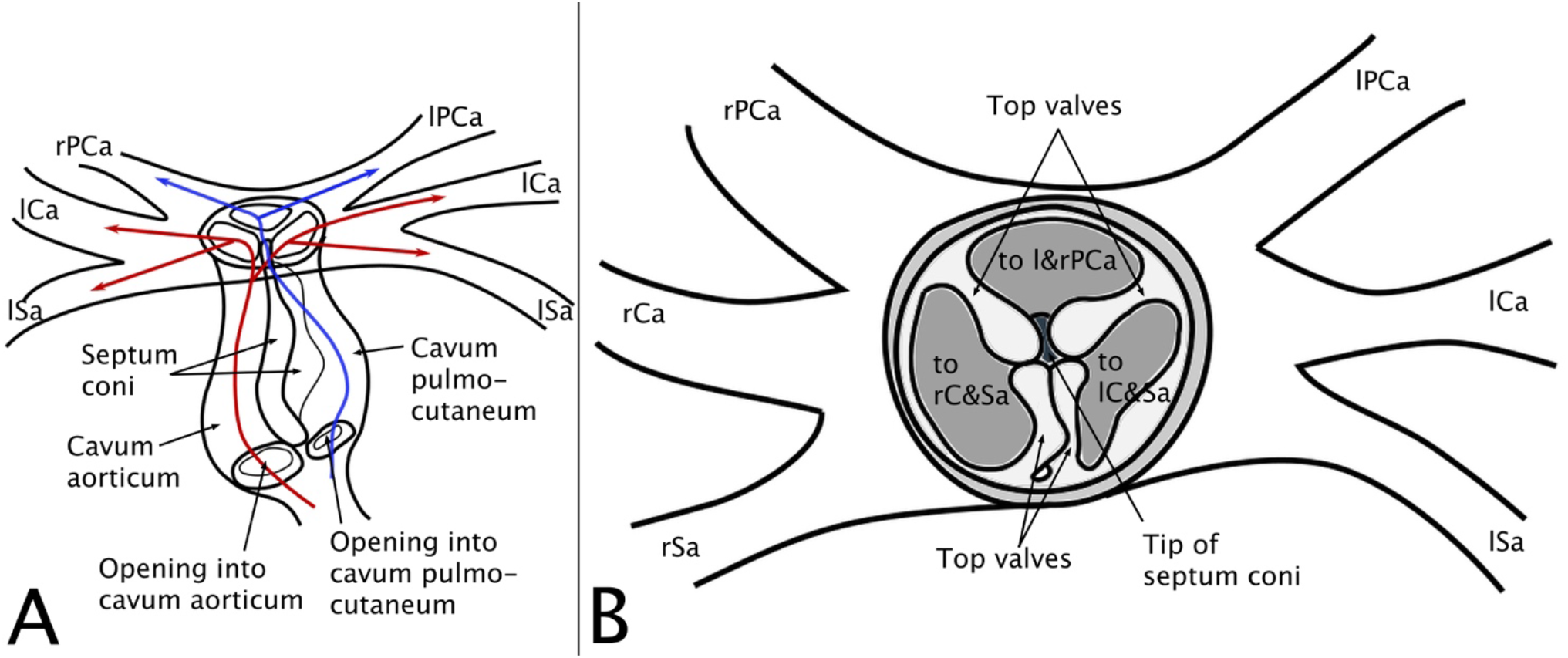
Schematic drawing of the potential separating mechanisms of the conus arteriosus as suggested by the morphology of a tadpole at Gosner stage 29. A: ventral view. B: rostral view. In this model, oxygenated (red) and deoxygenated (blue) blood separately enter the conus arteriosus via the two openings at the conal junction with the ventricle. The two streams are kept separated throughout the whole length of the conus by the septum coni, which divides the conus into the cavum aorticum and the cavum pulmocutaneum. The cavum aorticum then leads into the two more ventral openings of the conus into the aortic arches, the cavum pulmocutaneum reaches into most dorsal one. The Y-shaped separation forming those three openings at the top of the conus is formed by the two top valves, each consisting of two separated leaflets that contact the tip of the septum coni protruding into the middle of this opening. At atrium, Gi gills, lCa left carotid arch, lPa left pulmocutaneus arch, lSa left systemic arch, rCa right carotid arch, rPCa right pulmocutaneus arch, rSa right systemic arch, Ve ventricle.

After Gosner stage 25 (Fig. 10), conus development starts to slow down again. However, the thin sheets of tissue lining the conus walls, which connected top and bottom walls, as well as septum coni, grow thinner. Since this lining has already been thinner on the left side of the conus, it vanishes sooner than the lining on the right. This slow reduction of this conus wall lining continues up to Gosner stage 28, leaving only the tip of the septum coni in contacts with small bottom valve at the left, with all other structures within the conus looking like distinct units by now (see Fig. 11B). Additionally, the valves and the septum coni grow thinner, leading to the general heart morphology to look more delicate. At this point, the scans show fewer nuclei in these structures.

After Gosner stage 30, there are no further changes to either the overall heart morphology or the structures within the conus arteriosus. But the heart continues to grow, taking up proportionally more space within the thoracic cavity. Since the lungs come to lie between the bottom of the animals’ mouth and its heart, the heart is pushed down to a more ventral position. This pattern becomes even more pronounced as the lungs grow larger up to Gosner stage 46, when the larva has turned into a small froglet and metamorphosis is complete.

### 4.2. Gills and aortic arches

No gills are present at Gosner stage 18, but the blood vessels originating from the area of the truncus arteriosus already extends into the general area where gills will soon develop.

At the transition from Gosner stage 18 to 19, the carotid arch starts to protrude into the outside, followed by the systemic and pulmo-cutaneous arches, forming what already resembles very small gills (see Fig 15A).

At the point of reaching Gosner stage 19, a small tuft of external gills is present. Already, each of the three aortic arches originating from the heart each reaches into its corresponding gill arch. The hyoid arches are forming a cover for this tuft of gills (see Fig. 15B).

This situation remains the same up until Gosner stage 23 (see Fig. 15B&C). Here, the hyoid gill cover starts to grow and soon contacts the animals’ ventral side. While this skin folds continues to grow, it starts to cover the external gills, but it initially maintains a small opening on both sides out of which the gill tufts protrude (see Fig. 16).

Soon, within the same Gosner stage, an asymmetric pattern within the development of internal gills starts to appear. The skin fold continues growing over the animals’ right gill towards its ventral side while only the right gill tufts start to withdraw and become shorter, so that they become fully covered by this skin fold. On the left side however, the opening remains and the gill tuft continues to protrude. The left gill tuft eventually does withdraw into the gill cavity as well, and the remaining opening of the skin fold becomes smaller. This never closes completely but remains as the animals’ opercular spout starting from Gosner stage 25 (see Fig. 17).

Once the gills are situated fully within the now formed gill chambers, they become more finely branched, potentially facilitating gas exchange. This pattern continues up until metamorphic climax (see Fig. 14). Only at Gosner stage 46, once the animal is already a small froglet, the gills start to recede very rapidly and are completely gone at the end of what is considered larval development.

**Figure 13:**
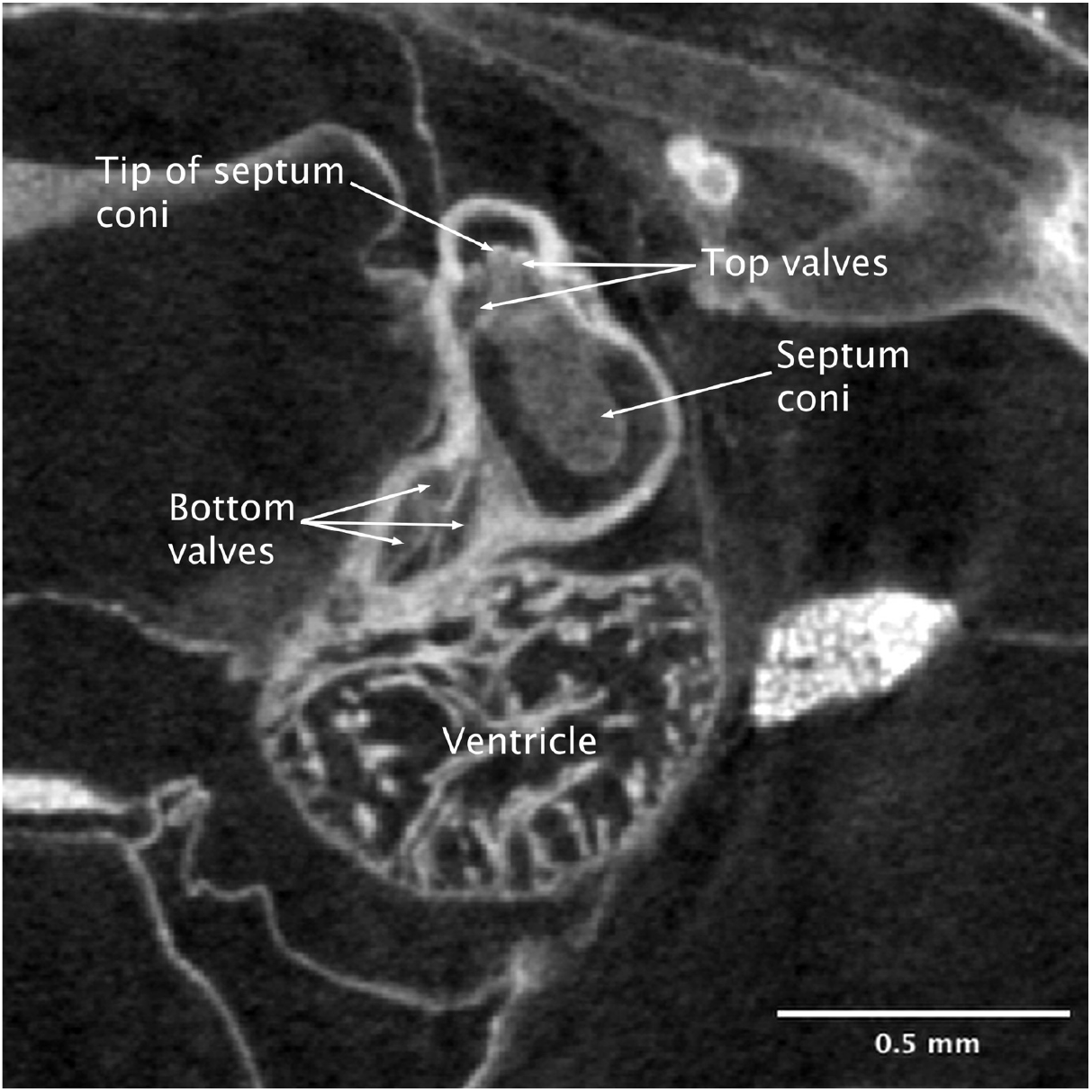
Virtual section through the heart of a tadpole at Gosner stage 40 showing the bottom valves (can be seen as either three separate ones or two main ones with one of them having an extra leaflet) which form two openings into the conus at the ventricular-conal juntion. Additionally, the septum coni and its tip reaching inbetween the top valves are shown.

**Figure 14:**
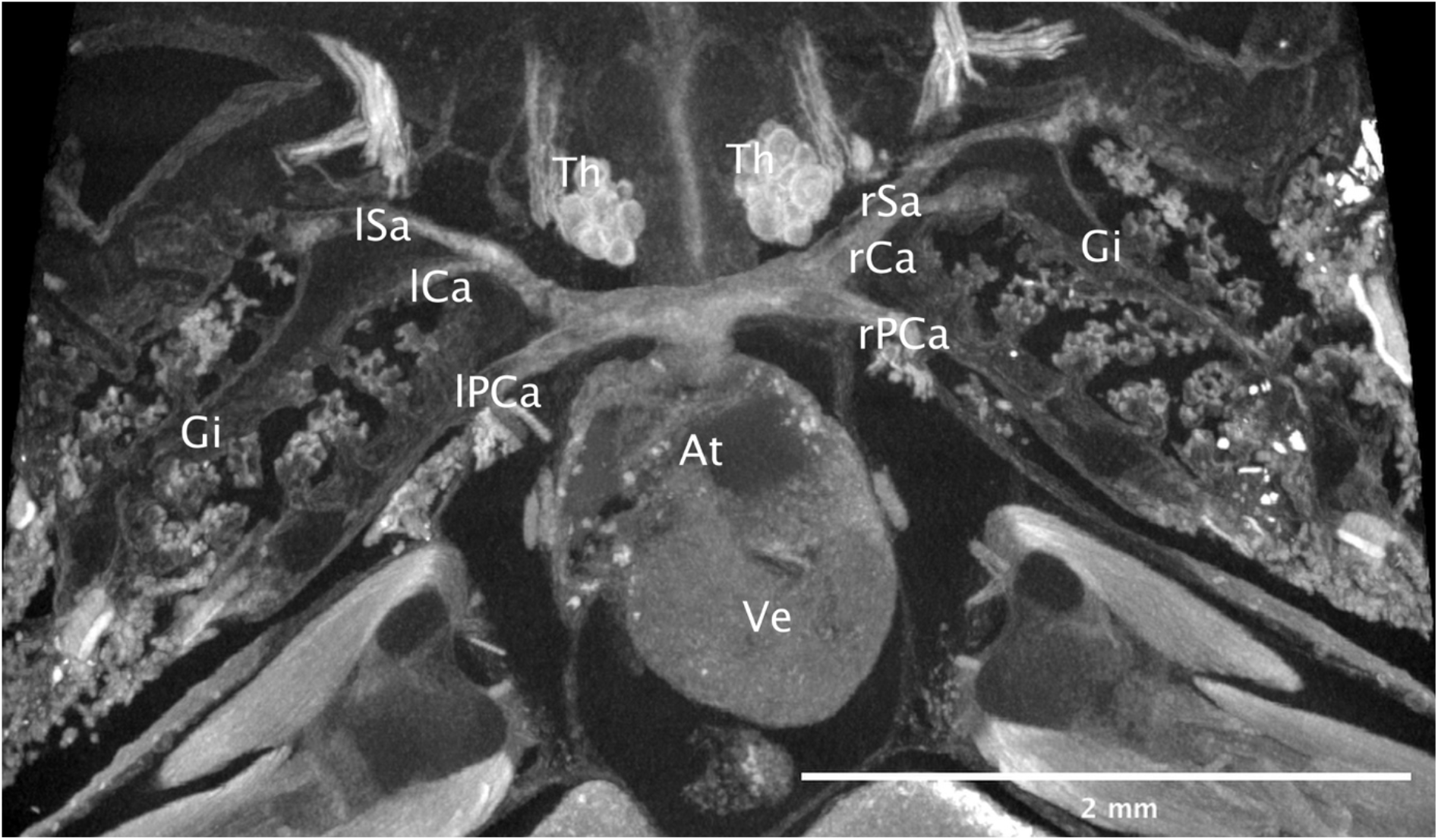
Gills and aortic arches at Gosner stage 43, almost at the end of metamorphosis. Dorsal view of a virtual cutaway. A small lung sack is already in place (not visible here). However, aortic arches still lead directly into the gills. At atrium, Gi gills, lCa left carotid arch, lPa left pulmocutaneus arch, lSa left systemic arch, rCa right carotid arch, rPCa right pulmocutaneus arch, rSa right systemic arch, Th thymus, Ve ventricle.

**Figure 15:**
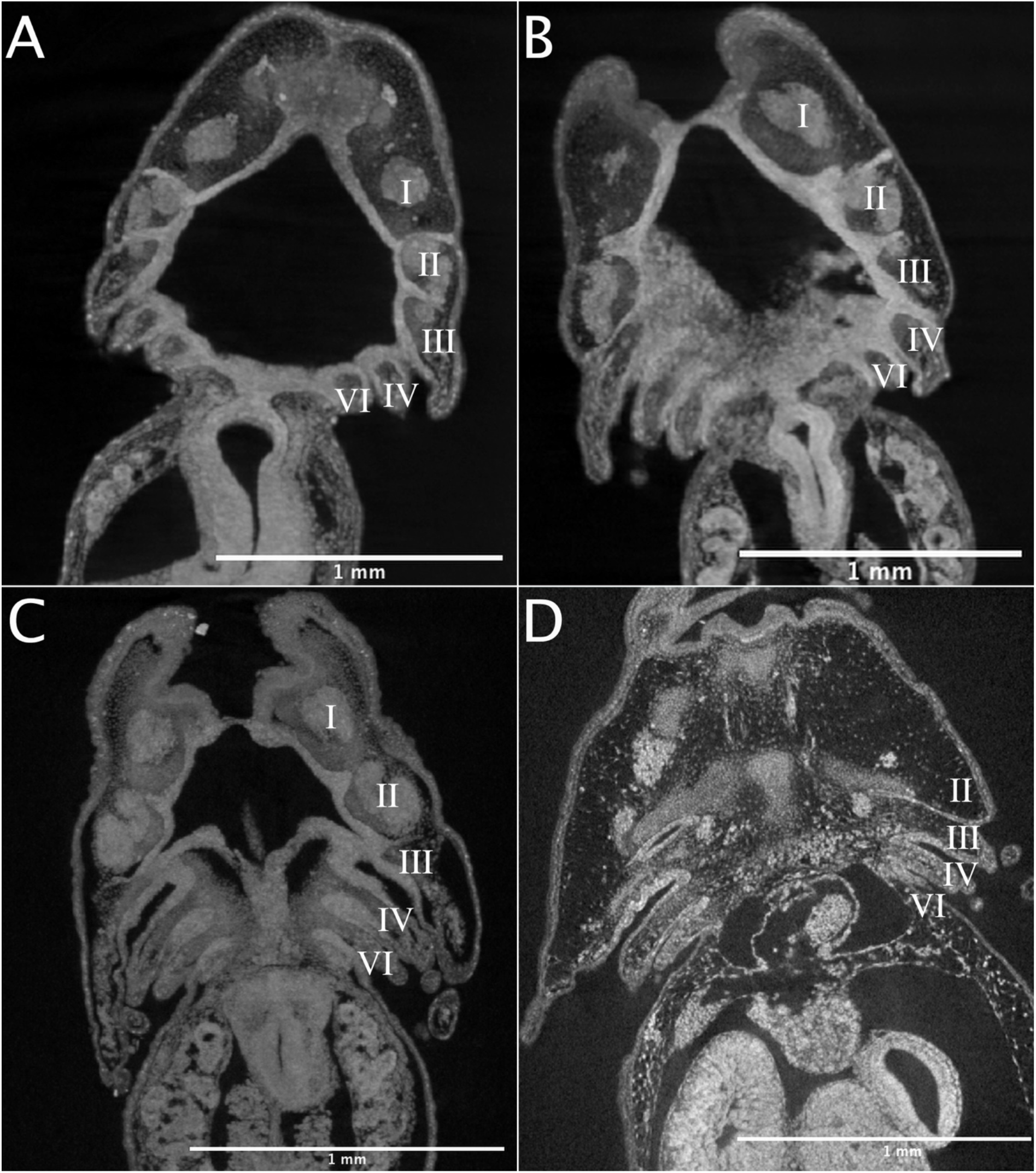
Overview over the development of the pharyngeal arches in virtual thick sections through the horizontal plane of tadpoles at different developmental stages. A: late Gosner stage 18. B: Gosner stage 19. C: Gosner stage 21. D: Gosner stage 23. I: mandibular arch. II: hyoid arch. III: third arch. IV: fourth arch. VI: sixth arch; Vth arch not present in anurans.

## 5. Discussion

This investigation began with the hope of demonstrating the correspondence between tadpole respiratory modes and cardiopulmonary development. We hypothesized that such a correlation between changing heart morphology and respiratory mode could illustrate how oxygenated and deoxygenated blood are organized within the anuran heart. Surprisingly however, the process of heart development seems almost unaffected by the breathing modes used.

Our results show that heart development is roughly finished at Gosner stage 28, even though lungs do not appear up until Gosner stage 43 (see Fig. 14), meaning that potentially separating structures within the conus arteriosus are already in place at a time when tadpoles still rely solely on gills and cutaneous breathing (see Fig. 10). This implies that, for most of their development, *Bufo bufo* tadpole hearts transport only deoxygenated blood that then gets oxygenated at the gills before it transports fresh oxygen to the rest of the body for most of their development, even though their heart morphology already shows the potential to function as an approximated double circulatory system.

This finding raises the need for an explanation for such seemingly counter-intuitive timing of heart development. A potential cause could be that the timing of heart development might be very conserved within anurans. So, while Bufonidae are known for only developing lungs at metamorphic climax and hence do not rely on an approximated double circulation up to this point, other species of frogs do (Nodzenski et al. 1989). Ranidae for example develop functional lungs much earlier in their larval phase than Bufonidae (Burggren 1989). *Xenopus laveis* starts to rely solely on its lungs very early in development and uses its gills only for filter feeding (Wassersug & Murphy 1987). So, while this developmental timing seems to make little sense for *Bufo bufo*, it could be essential for other anurans. However, so far there is very little developmental data on different anurans, so that such a conclusion cannot be made with great certainty. More studies on anuran, or even other amphibian, heart development would be needed, especially in a phylogenetic context.

Another possible explanation for the decoupling of tadpole respiratory modes from cardiopulmonary development could be that already separated streams of blood are necessary for correct heart development, even though both streams continue to contain only deoxygenated blood for most of development. This hypothesis and the one proposed for conserved heart development within anurans are not mutually exclusive, and they may even be two different facets of the same phenomenon. Several studies have already shown a strong influence of hemodynamic processes on heart development (e.g. D’Amato et al. 2016, Gitler et al. 2003, Moorman & Christoffels 2003, Person et al. 2005, Sedmera & McQuinn 2008). Those hemodynamic forces influence early heart development either by regulating how endocardial cells displace cardiac jelly as within the ventricle, or by regulating migration of endocardial cells into cardiac jelly, as within the outflow and the inflow tract (see Kirby (2007) and Rosenthal & Harvey (2010)). Those mechanisms play a crucial role in the formation of the atrial septum and heart valves, and therefore also the septum coni, which would mean that for the heart to be able to form blood-separating structures needed by the adult and later tadpole, streams of blood need to be separated early on. Such separated streams of blood in the early larval heart were actually observed by Jaffee (1963).

To our knowledge however, studies on anuran heart development have never considered the cardiac jelly. While it was noticed in 1920 as a “myoendocardial space” (Ingalls 1920) and was then first described as “cardiac jelly” by Davis in 1924, no authors working on anuran heart development seem to have taken notice of this. However, they did notice some of its peculiar properties. Ison (1968) for example noticed that precursors to the heart valves are already seen in the early embryo. Additionally, he observed that blood flow through the conus arteriosus causes the endothelium “to be thrown into folds” and he argued that the endothelium grows faster than the myocardium, leading to the formation of structures within the conus. Jaffee (1963) hypothesized that the two streams of blood within the developing anuran heart mold the atrial septum. He claimed that this tissue probably needs vascular forces for proper development because it shows far fewer nuclei than developing tissues normally do. While his hypothesis seems to be correct, the reasoning behind it seems to be flawed: the few nuclei he saw are probably cells of the endocardium migrating into the cardiac jelly and while vascular forces actually do seem to be essential for heart development, their effects are probably not only because of a lack of nuclei (see (Manasek 1970) and (Icardo & Fernandez-Terán 1987). Simons (1957) even showed the importance of blood flow on anuran heart development experimentally. By cutting the pulmonary vein in some specimens, he could show that the left atrium grew smaller than the one in intact specimens did. However, despite all the evidence showing the importance of vascular forces on vertebrate heart development, not much is known about the biomechanical properties of the specific tissues. Hence, it is too early draw firm conclusions from this phenomenon.

Nevertheless, further consideration of the cardiac jelly in embryonic heart development can aid in clearing up the enigmatic morphology of the structures within the anuran conus arteriosus. It does, for example, offer a partial explanation as for why previous researchers found differences in the number of valves both at the top and at the bottom of the conus, as well as differences in the spiraling pattern of the septum coni. Since all those structures can be traced back to one continuous tissue that once lined the whole conus, different structures like the valves are not such distinct units as one might assume when taking only the adult morphology into account. This, taken together with the different histological sectioning planes used by Ison (1967), can easily lead to such discrepancies regarding the morphology as are found in previous literature.

By taking into account developmental data and by using 3D micro-CT imaging, we were able to make a more detailed and comprehensive description. Our results show that the conus has two major valves and a small extra leaflet at its junction with the ventricle, as well as two valves at the top with two small leaflets each, which, together with an upward-protrusion of the septum coni, form a Y-pattern (see Fig. 11B and 12).

Additionally, starting from Gosner stage 23, we found a small extra opening at the junction between the ventricle and the conus, separate from the large one that is surrounded by the row of valves. Previously, de Graaf (1957) showed that the septum coni divides the conus into two chambers, the cavum aorticum that lies more ventrally and is in contact with the main orifice at the junction between ventricle and conus, and the cavum pulmocutaneum lying more dorsally (see Fig. 11B and 12). However, de Graaf also mentioned that it looks like the cavum pulmocutaneum has no direct communication with the ventricle, while also pointing out that this seems unlikely. We were able to find such a postulated opening connecting the cavum pulmocutaneum with the ventricle. It is, however, a rather small and narrow gap lying caudal to the main orifice, making it rather easy to miss when relying on histological sections alone. This finding would fit well with the model of blood separation of the Classical Theory, which describes the ventricular systole as initially sending less oxygenated blood into the cavum pulmocutaneum. Then, at a later point within the ventricular systole, the conus arteriosus is said to contract as well, causing the septum coni to contact with the conus wall at the base of the orifice at the junction between ventricle and conus, so more oxygenated blood can then enter the cavum aorticum (see de Graaf 1957 for a detailed review on this). Thus, it may be that this small extra opening leading into the cavum pulmocutaneum gets closed off by the septum coni during the first ventricular systole. The theory for how the ventricle keeps blood separated in the first place is via a pressure gradient that directly draws deoxygenated blood from the right atrium into the cavum pulmocutaneum, while the more-oxygenated blood is held back within the trabeculae of the ventricle, before it gets drawn into the cavum aorticum during a second part of the ventricular systole. However, those mechanisms need to be studied further since most of this theory is based on theoretical considerations and extrapolation from the similar lungfish heart that, unlike anurans, has a septum in its ventricle instead of trabeculae (Johansen & Hanson 1968).

Another aspect to be emphasized about the small extra opening we found at the base of the conus is its developmental timing. It first starts to appear at Gosner stage 23, when there is still only one cavum within the conus where blood could potentially flow (see Fig. 11A). But during Gosner stages 24 and 25, the right side of the conus, which aligns with this small opening, starts to form an additional cavum, and together with the valves from the left side of the conus and the upwards protrusion of the septum coni, forms the Y-shaped structure found at the top of the conus (see Fig 11B and 12). This timing in development, taken together with the known properties of cardiac jelly, suggest that this extra opening enables a new blood-flow pathway through the conus that in turn starts to “carve out” its own cavum, contributing significantly to the formation the distinct structures found within the adult anuran conus arteriosus.

Following the two cava of the conus arteriosus up to the top rows of valves shows that blood with different oxygen content could potentially be kept separated along the whole length of the conus by the spiraling of the septum coni (see Fig. 6). Then, at the top of the conus, the upwards protrusion of the septum and the two valves forms three chambers. The two more rostral ones are in contact with the cavum aorticum and lead into the carotid and systemic aortic arches. The caudal-most opening is in contact with the cavum pulmocutaneum and leads into the pulmo-cutaneous aortic arch (see Fig. 12). While morphology alone clearly cannot prove whether these structures actually carry out the proposed functions, our results show that anuran heart morphology would allow an approximated double circulation, consistent with the considerations of the Classical Theory. Our descriptions of heart morphology can provide a strong basis for future research focusing more on the function of the anuran heart.

The relative timing of heart development and formation of the gills and aortic arches is clearly important in the quest to understand anuran heart function, but much about those structures remains unknown. One important aspect is the transition from external to internal gills during larval development. Our results show that the external gills start to withdraw into what will become the gill chamber owing to a skin fold that starts from the hyoid and carotid arches and grows towards the ventral side and finally fully covers the animals right gills but leaves open a small opercular spout on the animal’s left side (see Fig. 16 and 17).

**Figure 16:**
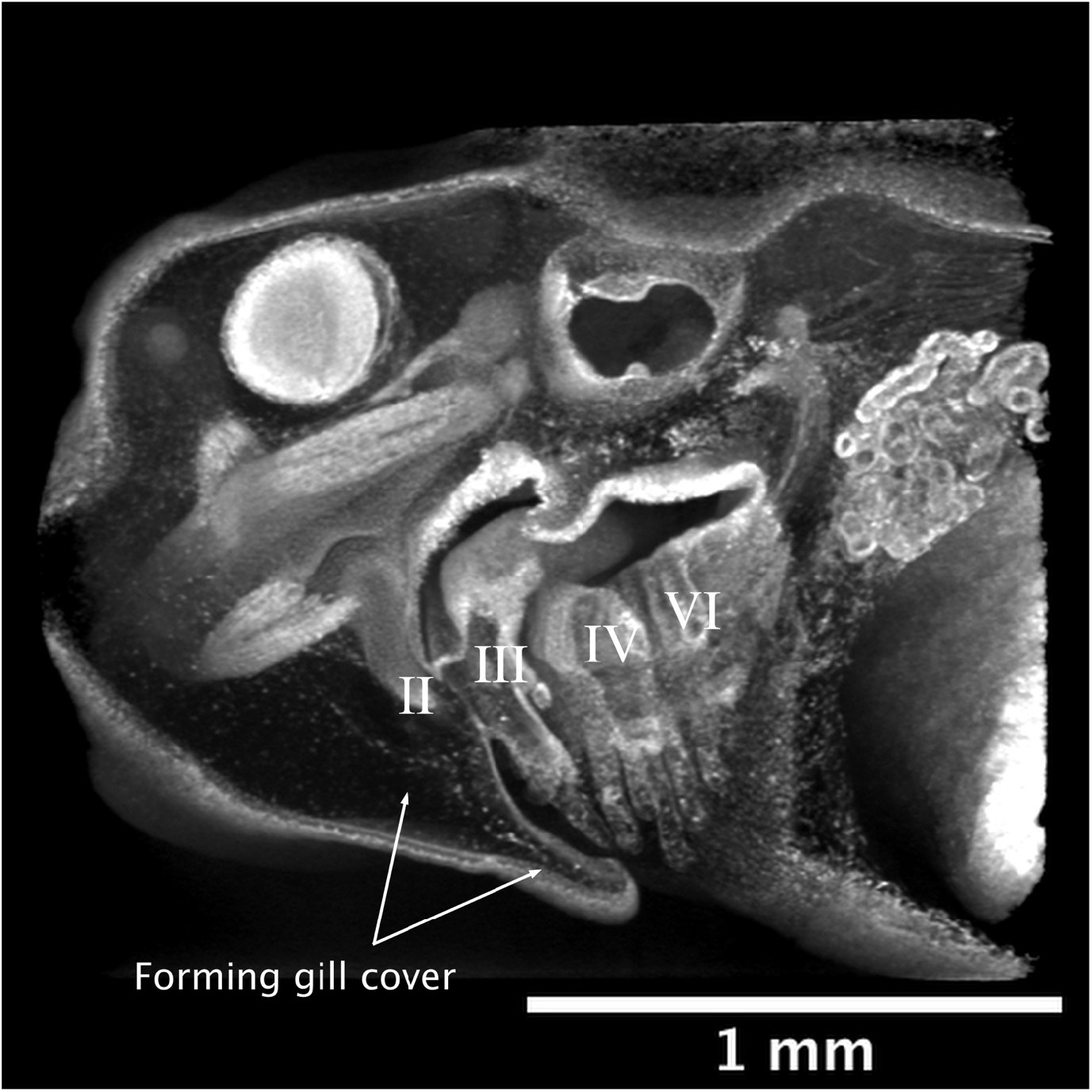
Virtual sagittal cutaway of a tadpole at Gosner stage 23 in lateral view showing the gill cover forming. II: hyoid arch. III: third arch. IV: fourth arch. VI: sixth arch.

**Figure 17:**
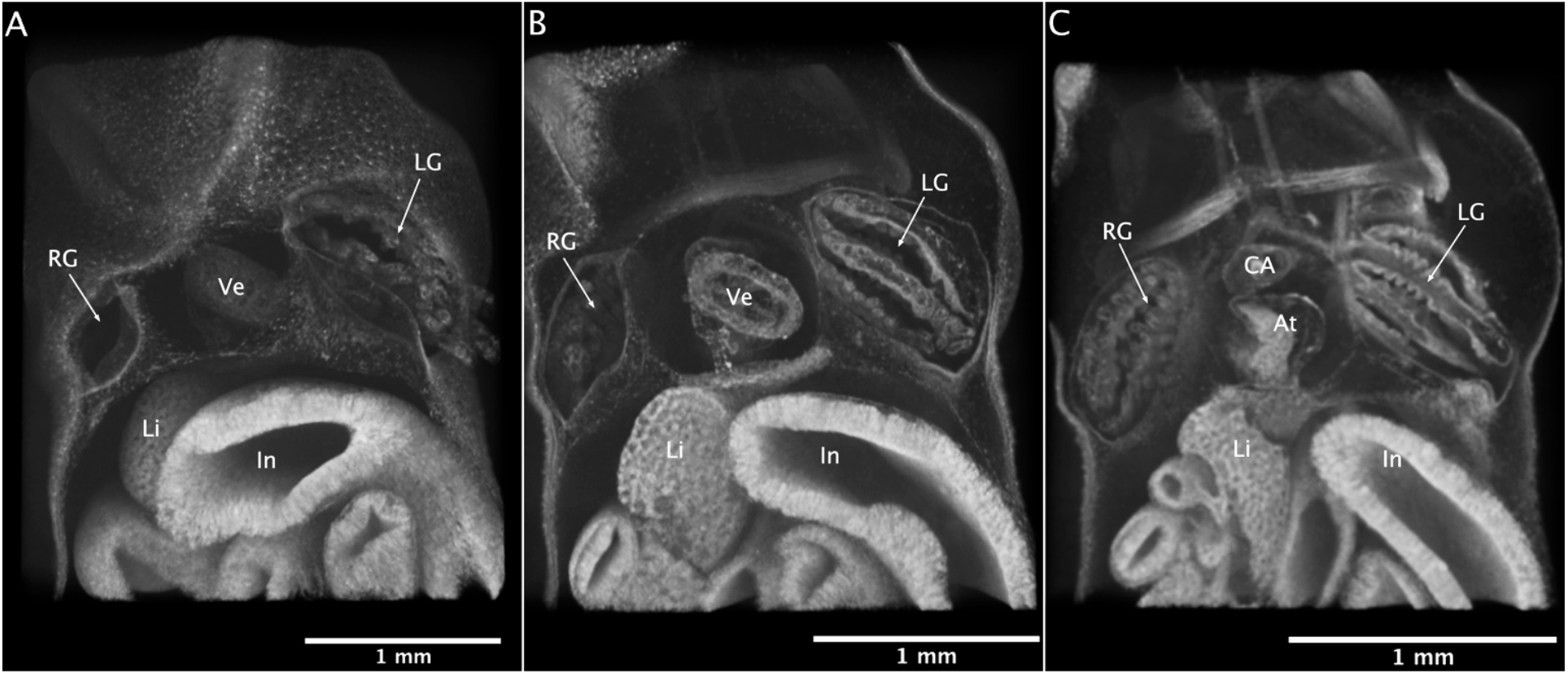
Virtual horizontal cutaway of the ventral side of a tadpole at Gosner stage 23 showing the asymmetry of the process of gill absorption and covering, and formation of the opercular spout. At atrium, CA conus arteriosus, In intestines, LG left gills, Li liver, RG right gills, Ve ventricle

Our observation of this transition from external to internal gills raises the question of why it even takes place at all. Curiously, salamander larvae do not show this pattern of replacing their simple external gills by morphologically more complex internal gills (McIndoe & Smith 1984), but rather their external tuft of gills simply grows bigger and becomes more vascularized (Guimond & Hutchison 1976, Malvin 1989). However, despite being clearly peculiar, to our knowledge there are no studies or hypotheses as to why this should be the case. One possibility based on what we know at the present is that it is simply a consequence of the general larval body shapes of those different amphibian orders. This idea is based on our observation that around the time anuran tadpoles start to change from external to internal gills, their body shape starts to change from a slender and elongate early tadpole to a very plump and round animal with a tail, while salamander larvae remain slender during their whole larval development. Hence, it could simply be that the shape tadpoles need to have in order to develop into an adult frog has an unfavorable ratio of surface to volume, which is relevant, because the skin is an important area of gas exchange (Burggren & West 1982). Thus, in order to compensate for that, anurans started to develop more efficient gills that enable more gas exchange. Since it seems that no research has been done on this phenomenon, this hypothesis serves as a reasonable starting point based on the limited data that we have.

As we were able to show, *Bufo bufo* develops its lungs only at metamorphic climax, as is typical for bufonids, whereas ranids and others usually develop lungs much earlier (Burggren & West 1982, Nodzenski et al. 1989, Wassersug & Murphy 1987). Geographical distribution is interesting in this context: Bufonidae occur much further north than many other anuran families (Metcalf 1923), so they likely adapted to development in colder climates and hence colder waters. This is significant because colder water generally holds more dissolved oxygen (Burggren & West 1982), meaning that a lack of oxygen in their respiratory medium is not a likely environmental pressure. On the other hand, anurans native to warmer climates might need to have at least the possibility to use the lungs earlier in case of high temperatures significantly decreasing the available oxygen in water. It would be very interesting to see whether this ecological pattern could hold true with more phylogenetically relevant data of anuran respiratory development; however, significantly more phylogenetically relevant development data will be needed to get a clearer picture of this phenomenon.

Generally, it has to be said that phylogeny and ecology have been hardly considered when trying to resolve the enigma that is anuran respiration and heart function. So, while it seems reasonable that mixed, semi-oxygenated blood is efficient enough for primarily aquatic anurans, which can compensate for their lack of a double circulation via cutaneous respiration, this seems more questionable, when, for example, looking again at Bufonidae. It seems rather unlikely that their thick skin enables them much cutaneous respiration despite being significantly drier than many other anuran skins. This, again, emphasizes the point that there is no one amphibian heart (Burggren 1988) and that for understanding both anuran respiration and heart function better, more data is needed.

The aim of this study was to offer a detailed description of the 3D morphology of a representative anuran conus arteriosus throughout larval development, as well as a description of gill and lung development in *Bufo bufo*. Our morphological data fit well with the model of heart function as described by the Classical Theory of functional blood separation in cardiac outflow. Further advances toward understanding anuran pulmo-circulatory function and physiology will require *in vivo* studies of blood flow through the anuran heart, but such data could never be usefully interpreted without a clear morphological understanding of the conus structure encompassing the relevant developmental stages and providing a comprehensive visualization of the anatomy. Additionally, we were able to describe how tadpoles transition from external to internal gills and we raise important questions about ecological factors when considering anuran heart development that are still significantly understudied.

## 6. Acknowledgments

The authors are grateful to Dr. Silke Schweiger and Georg Gasser (NHM Vienna) for providing specimens and for helpful advice and discussions, and to Prof. Mihaela Pavlicev and the Theoretical Biology Unit and Dept. of Evolutionary Biology for supporting this project as part of NK’s master’s work.

